# SAMson: an automated brain extraction tool for rodents using SAM

**DOI:** 10.1101/2024.03.07.583982

**Authors:** Daniel Panadero Soler, Mohamed Kotb Selim, Patricia Martínez-Tazo, Emma Muñoz-Moreno, Pedro Ramos-Cabrer, Pilar López-Larrubia, Silvia De Santis, Santiago Canals, Antonio Pertusa

## Abstract

Accurate brain extraction is a critical step in the analysis of rodent head magnetic resonance imaging (MRI) data. However, current methods often encounter difficulties in handling the diverse range of imaging setups, resolutions, and experimental conditions that are commonly found in this field. Based on the Segment Anything Model (SAM), we introduce here SAMson (SAM for Segmentation Of Neuroimages), an automated tool for robust rodent brain extraction. SAMson integrates a bounding box generator and a mask prediction pipeline, offering fully automated and semi-automated modes to address varying experimental complexities. The performance of SAMson was evaluated using three multi-centre rodent MRI datasets annotated at the pixel level, which differed in terms of acquisition parameters, resolution, and animal age groups. SAMson demonstrates superior performance to existing methods, including BET, RBM, and BEN, in terms of segmentation accuracy, with Jaccard indices exceeding 90% across datasets. The semi-automated mode demonstrates particular efficacy in challenging scenarios, including low-resolution images and cases requiring refined mask precision. In contrast to conventional volumetric techniques, SAMson identifies errors at the level of individual slices, thereby enabling rapid and targeted correction when needed. By providing open-source access, SAMson aims to support large-scale research workflows and advance translational neuroscience. The curated data can be downloaded from https://doi.org/10.20350/digitalCSIC/17000, and the code is available at https://github.com/CanalsLab/SAMson.

## 1 Introduction

Magnetic resonance imaging (MRI) is a fundamental technique in preclinical neuroscience that provides functional and structural information about the brain. This non-invasive and longitudinal method has demonstrated significant translational validity, as preclinical observations in animal models can be directly compared with human observations [1,2].

Brain extraction, or skull stripping, is the segmentation of brain tissue from the skull and other tissues in head MR images. This process, typically performed at the outset of preprocessing pipelines, relies on masks outlining the region of interest (ROI) [3]. Segmentation errors can compromise the reliability of scientific observations and the reproducibility of the results [4]. Consequently, researchers often resort to manual extraction, a labor-intensive process incompatible with the growing demand for large datasets to enhance scientific inquiry. Numerous algorithms have been developed to automate brain extraction [5]. Automation not only streamlines workflows but also reduces inter- and intra-operator variability, ensuring robust performance across datasets from different facilities and experimental conditions [3].

The current gold standard in automatic skull stripping is the Brain Extraction Tool (BET) offered by the FMRIB Software Library (FSL) [6]. However, the advent of deep learning methods has introduced several medical image segmentation tools, with U-Net-based models [7] achieving widespread adoption [8]. While effective for human brain segmentation, these methods often fail to adapt to rodent MRI due to the diversity of imaging setups and parameters [9]. An additional challenge is the lower image resolution relative to the brain-skull distance in rodents, complicating brain extraction [10]. Consequently, extensive manual editing is frequently required, rendering the process nearly as laborious as manual mask extraction itself.

A pioneering work in promptable image segmentation is the Segment Anything Model (SAM) [11], built on Vision Transformer (ViT) architecture [12] and trained on the SA-1B dataset (1.1 billion masks from 11 million images). This extensive training grants SAM robust zero-shot generalization capabilities [13], often outperforming fully supervised extraction methods across diverse segmentation tasks without retraining or fine-tuning [14]. However, the training dataset of SAM primarily consists of natural images, which differ significantly from medical images [15]. Natural images typically feature color encoding, well-defined object boundaries, and a clear separation between foreground and background, with a relatively balanced size distribution. In contrast, most medical images are grayscale, with ambiguous and complex object boundaries, low contrast, and overlapping characteristics between foreground and background [14,16,17]. Not surprisingly, studies show that SAM generally performs worse in medical image segmentation (MIS) compared to natural images [14,18,19,20].

Despite this limitation, SAM has demonstrated superior performance compared to other interactive MIS methods [21] and, in some cases, delivered zero-shot results surpassing state-of-the-art tools [13]. For MRI applications, SAM consistently segments medical images across various modalities and aspect ratios [17]. Specifically, in brain extraction from human MR images, SAM outperformed BET across a wide range of imaging modalities, contrasts, and pathologies [22]. Previous studies have evaluated the zero-shot application of SAM to medical imaging [13,14,15,17,18,19,21,22], often using Ground-Truth (GT) masks to create prompts, referred to as “oracle performance”. Other efforts focused on optimizing the manual input of prompts [23] or adapting SAM for the segmentation of medical images [16,20,24]. However, limited attention has been directed toward the critical aspect of automation. Due to its prompt-based design, developing automated pipelines based on SAM presents a challenge [20]. Nonetheless, a model trained for promptable segmentation can serve different purposes when integrated into a larger algorithmic system [11].

Building on the above strategies, we present SAMson (SAM for Segmentation Of Neuroimages), an automated tool for brain extraction from rodent MR images. SAMson integrates an independent generator of object-bounding boxes as prompts for SAM, enabling the automatic production of accurate masks. We validated SAMson on multi-center preclinical MRI datasets from mice, demonstrating operator-independent segmentation results closely matching manually created masks by an expert. This advancement streamlines preclinical research workflows and supports translational studies. More broadly, SAMson establishes a foundation for SAM-based automated segmentation pipelines, with potential applications across diverse medical imaging tasks.

## 2 Methodology

### 2.1 Datasets

Three distinct MRI datasets have been acquired and curated to evaluate SAMson’s performance (Table 1) across varying acquisition protocols, magnetic field strengths, and image resolutions, thereby testing its robustness and transferability in diverse preclinical neuroimaging settings. The primary dataset (D1) comprises a cohort of 53 mice, stratified into four different age groups: 16 subjects at 3 months of age (m.o.a.), 11 at 4.5 m.o.a., 11 at 6 m.o.a., and 15 at 12 m.o.a. To assess the generalizability of SAMson, we employed two additional datasets acquired in different MRI facilities and composed of 3 m.o.a. mice: D2, which encompasses 10 subjects, and D3, including 12 subjects.

**Table 1:** Acquisition parameters of the three datasets used for evaluation. D1: Images acquired on a 7-T scanner (Bruker, BioSpec 70/30, Ettlingen, Germany) using a T2-weighted RARE (Rapid Acquisition with Relaxation Enhancement) sequence. D2: T2-weighted sequence on a 7-T scanner (Bruker, BioSpec 70/30, Ettlingen, Germany) equipped with an actively shielded gradient system. D3: 11.4-T MRI scanner (Bruker BioSpec USR 117/16, Ettlingen, Germany) using a T2-weighted RARE sequence.

| Dataset | TR / TE (ms) | FOV (mm) | Matrix Size | Resolution (mm) |
| --- | --- | --- | --- | --- |
| D1 | 3000 / 7.698 | $18 \times 15$ | $108 \times 90 \times 16$ | $0.1667 \times 0.1667 \times 0.8$ |
| D2 | 2500 / 33 | $34 \times 34$ | $256 \times 256 \times 22$ | $0.1302 \times 0.1302 \times 0.8$ |
| D3 | 5500 / 26 | $17.5 \times 17.5$ | $175 \times 175 \times 48$ | $0.1 \times 0.1 \times 0.3$ |

All D2 images were either utilized in two previously published studies [25,26] or unpublished (authors acknowledged), but were not publicly released. In our work, all images in the three datasets have been manually annotated for brain segmentation and released.

### 2.2 Pipeline overview

SAMson’s architecture consists of two primary components: an object detector and a mask predictor. In brief, the object detector produces the necessary prompts that are used by SAM to segment the brain accurately. In this section, we describe the methodology in detail, following the steps in Fig. 1.

**Fig. 1:**
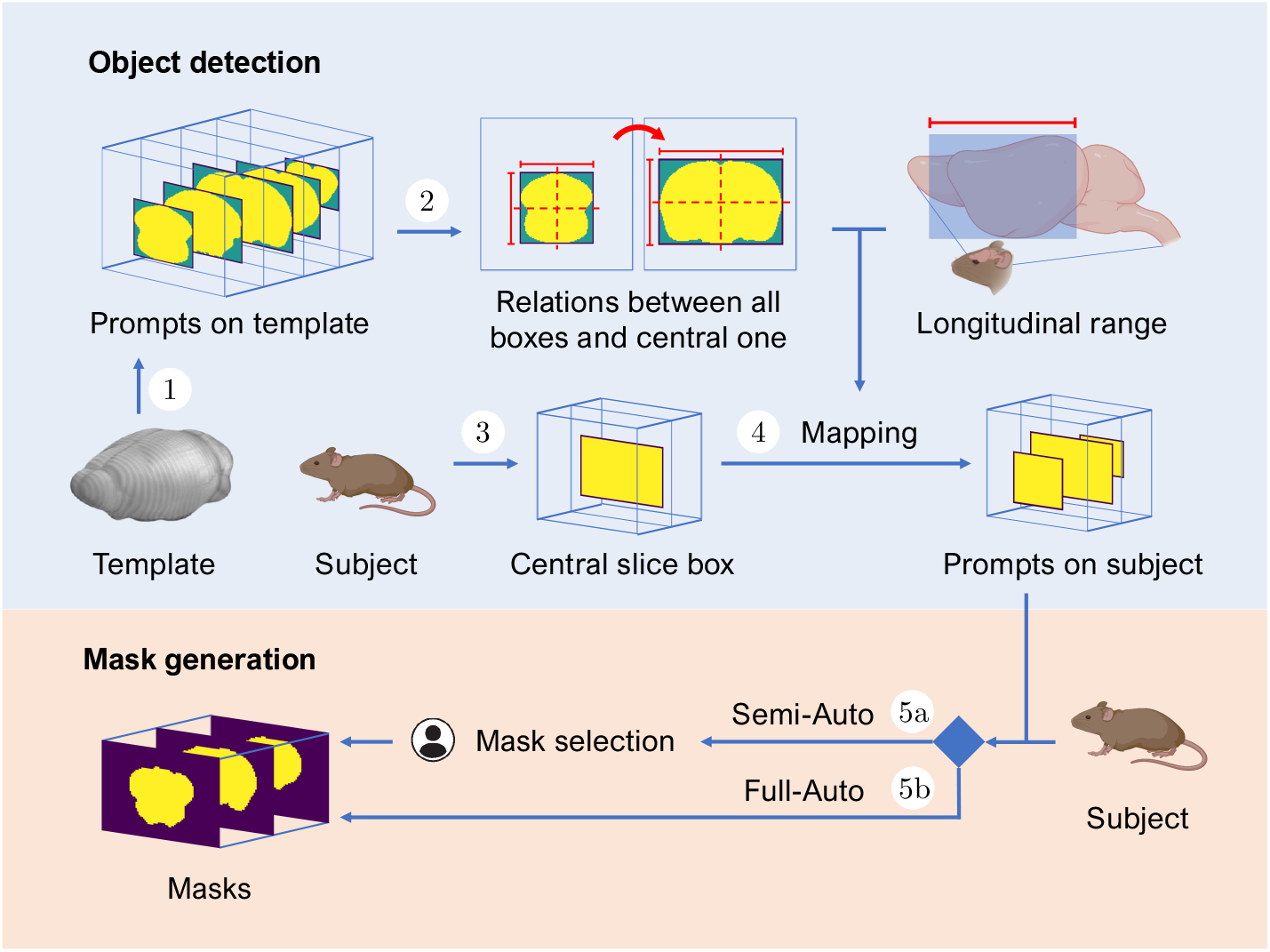
SAMson pipeline. 1. Template prompt generation; 2. Calculation of relative coordinates; 3. Definition of the central slice box per subject; 4. Mapping; 5a. Semi-Auto mode; 5b. Full-Auto mode.

### 2.3 Object detection

To generate input prompts, SAMson utilizes bounding box mode, which outperforms point prompts by enhancing prediction stability. Bounding boxes offer a less ambiguous spatial context for the ROI, capturing essential features such as the object’s size, location, and intensity profile [13,16,17,19,20,21]. Since box-based prompting is sensitive to slight offsets of the bounding boxes [13,16,17], the precise definition of bounding box coordinates is the first and most critical step for accurate automatic segmentation in SAMson (see Algorithm 1).

1. **Template prompt generation**. Using a mouse brain-extracted template [27] integrated into the tool, SAMson automatically generates bounding boxes enclosing the brain region in each slice. In this approach, bounding boxes are aligned with the average anatomical space of the mouse brain, reducing the task of automated prompting to a registration operation.
2. **Calculation of relative coordinates**. From the template prompts generated in (1), we extract the relative coordinates of all boxes with respect to the central slice box in the template space.
3. **Definition of the central slice box per subject**. Two primary factors contribute to variability in locating the ROI: the antero-posterior coverage of the image set (field of view, FOV) and the ROI size and location within the FOV. By default, SAMson sets the FOV to the typical range in rodent MRI, spanning from the anterior tip of the frontal cortex to the posterior occipital cortex. Users can specify custom coordinates for different ranges to adjust to the FOV, just by specifying the most anterior and posterior slices in their experiment. To address variability in size and location of the brain relative to the FOV, SAMson employs a brain segmentation function based on Otsu’s method [28], implemented in the DIPY (Diffusion Imaging in Python) library [29]. This method is most reliable on central slices and is used to generate a bounding box for the subject’s central slice. This central slice box serves as the basis for mapping and generating subsequent box prompts, compensating for inter-individual variability and differences in head positioning within the FOV.
4. **Mapping**. The relative coordinates are transposed to the subject space through linear interpolation, with flexibility for any number of slices and antero-posterior ranges. These coordinates are then applied to the previously generated central slice box to produce the complete set of bounding boxes in the subject space.

### 2.4 Mask generation

We adapted SAM’s testing pipeline^1^ to accept the generated prompts and perform iterative mask prediction across consecutive slices. We implemented two different mask generation modes, Semi-Auto and Full-Auto, differing in the level of operator dependence.

5a. **Semi-Auto mode** (see Algorithm 2). It leverages SAM’s capability to make three mask predictions for a single prompt, intended for whole object, part and subpart segmentation [11,15]. In our case, these correspond to varying levels of brain extraction, increasing the odds of producing a suitable mask.

5b. **Full-Auto mode** (see Algorithm 3). The mask predictor is configured to produce one single output per slice, automatically assembling all the results into a corresponding volumetric mask.

SAMson uses the largest SAM model, ViT-H (with 32 transformer layers and 636M parameters), as it is the default option and has been shown to provide substantial performance improvements over smaller models in MIS [17].

### 2.5 Mask correction

All generated masks, including the one produced using Otsu’s method, are refined through morphological image processing to eliminate outgrowths marginally connected to the ROI and isolated false-positive voxels (see Algorithm 4). For outgrowth removal, the mask is eroded to detach the undesired lobes, which are then dilated and subtracted from the original mask. Since dilation and erosion are not strictly inverse operations, residual voxels may persist along with small disconnected components hallucinated by SAM [11]. SAMson automatically removes these errors.

## 3 Results

The quality of the masks generated using SAMson was compared with GT masks extracted by an expert researcher. For this quantitative performance assessment, we used the Jaccard index metric [30], which evaluates the overlap between predicted and GT masks. Ranging from [0, 1], a higher value indicates a larger similarity between reference and predicted masks.

### 3.1 SAMson performance analysis

Both the Semi-Auto and Full-Auto SAMson versions yielded highly accurate segmentation results (Table 2), showing excellent agreement with manually extracted masks. As expected, the Semi-Auto method outperformed the Full-Auto, but the differences were small, confirming the strong performance of the automated approach. The results were consistent across all three datasets (D1–D3) and ages, with slightly lower performance observed for older animals (Fig. 2-a,c). Changes in the imaging protocol or number of image slices had no impact on performance and did not require code adjustments.

**Table 2:** Average Jaccard index results in percentage with their standard deviation. Statistical comparison between Full-Auto and Semi-Auto results using paired t-test: D1, *t*(104) = 6.5, *p* < 0.001; D2, *t*(18) = 3.1, *p* < 0.01; D3, *t*(22) = 3.2, *p* < 0.01.

| Method | D1 | D2 | D3 |
| --- | --- | --- | --- |
| SAMson Full-Auto | 90.12 ± 2.47 | 95.20 ± 0.74 | 95.52 ± 0.46 |
| SAMson Semi-Auto | **92.74 ± 1.58** | **96.12 ± 0.47** | **96.15 ± 0.46** |

**Fig. 2:**
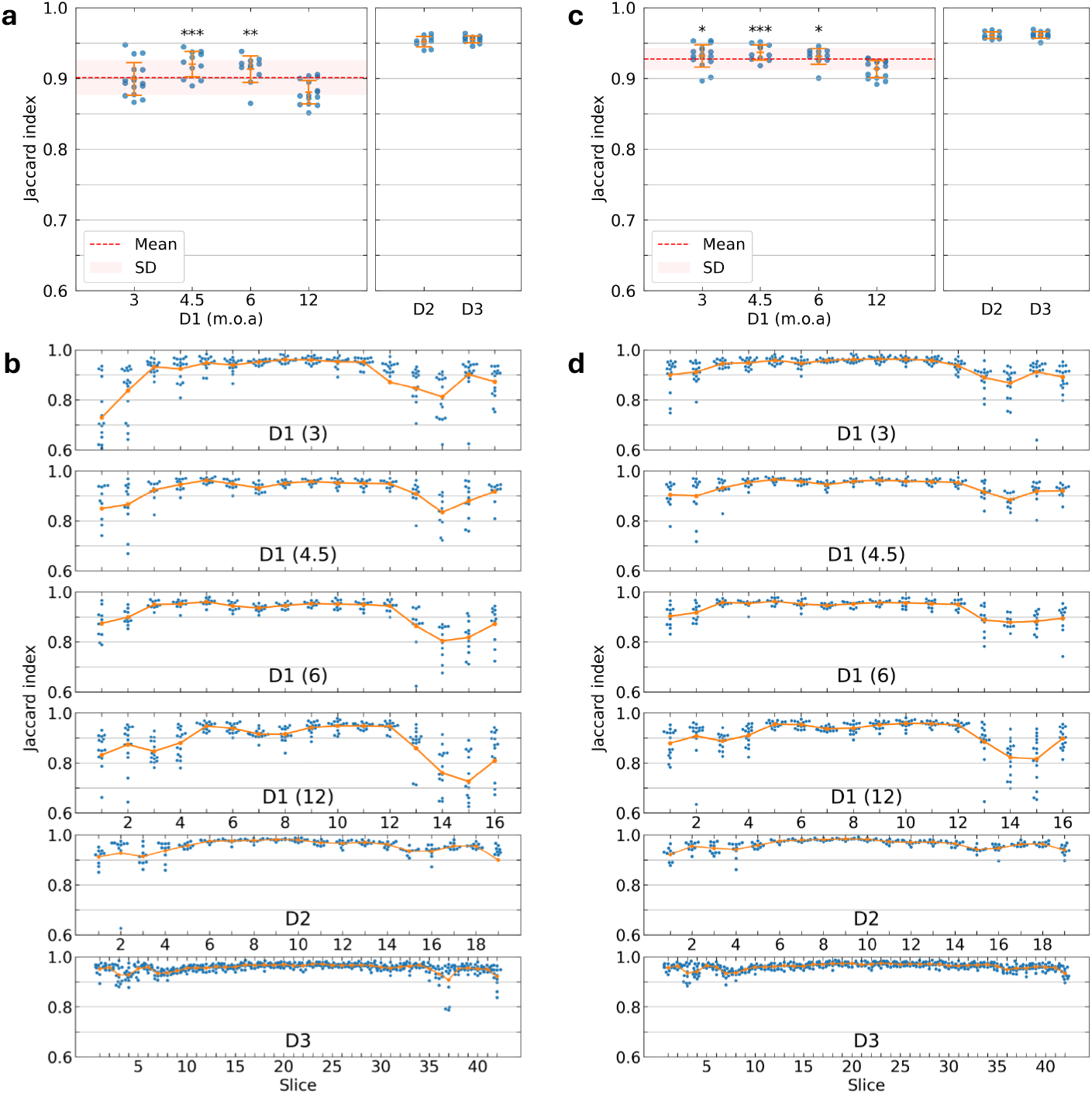
Brain extraction performance of SAMson in Full-Auto (a-b) and Semi-Auto (c-d) modes. Jaccard index computed relative to manual masks defined by an expert. Subjects from D1 were categorized by age group (m.o.a.: months of age). For D2 subjects, the first and last 2 slices, and for D3, the first 4 and last 2 slices were excluded from the evaluation due to the absence of brain tissue. In total, the analysis included 848 generated masks from D1, 190 from D2, and 504 from D3. a, c) Blue dots represent the average performance for each subject, computed across all slices, with the mean and SD in orange. For D1, global mean and SD across age groups is shown in red. One-way ANOVA showed a significant age effect in (a) (*f* (4, 52) = 9.9, *p* = 3·10^−5^) and (c) (*f* (4, 52) = 8.1, *p* = 1.7 · 10^−4^). Post-hoc analysis with unpaired t-test demonstrated the lower performance in the 12 m.o.a. group (Holm adjusted p-values vs. 12 m.o.a.: *p* < 0.05 (∗), *p* < 0.01 (∗∗), *p* < 0.001 (∗∗∗). b, d) Each data point represents the performance score for each slice in the antero-posterior axis. Orange lines connect the mean values for each slice group within their respective dataset or age group.

Segmentation performance was moderately influenced by image resolution (D1<D2<D3) (Table 2), and the advantage of the Semi-Auto method diminished with higher-resolution images (improvements of 2.9%, 1%, and 0.6% for D1, D2, and D3, respectively; see further analysis below), aligning with a trade-off between time spent on increasing resolution and preprocessing.

Fig. 3 presents representative examples of three typical cases (one per dataset) using the Full-Auto method. Overall, these findings demonstrate the tool’s robustness across varying experimental conditions, operators, and MRI facilities.

**Fig. 3:**
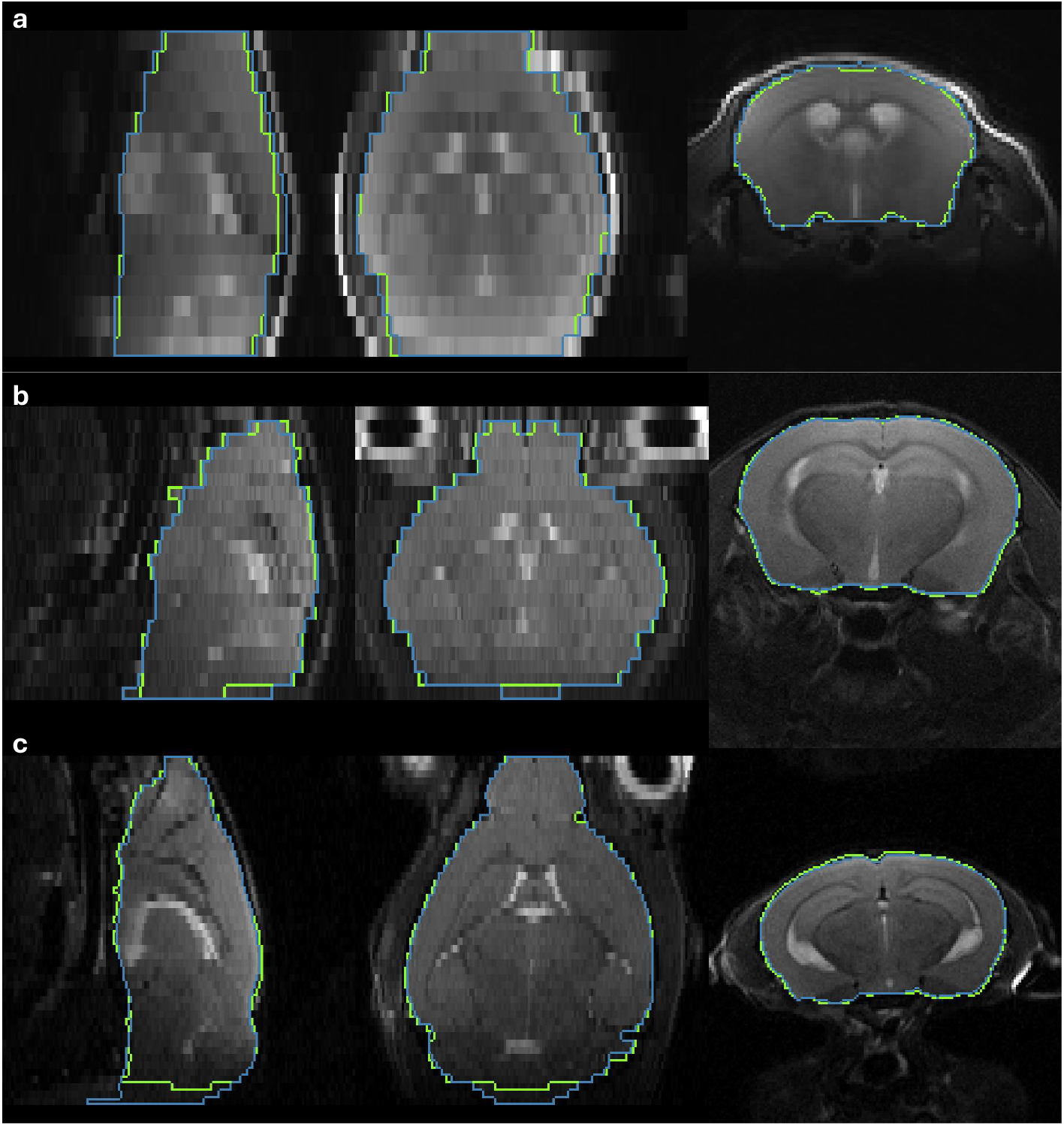
Representative brain masks extracted with SAMson. Green outlines indicate manually drawn brain masks; blue outlines represent automatically generated brain masks. Panels show one typical case from each dataset: a) D1, b) D2, and c) D3; with average Jaccard results: 0.9023, 0.9528, and 0.9568, respectively.

We further evaluated performance along the antero-posterior axis, across slices (Fig. 2-b,d), providing a more accurate depiction of the tool’s segmentation capabilities compared to averaged results. We found that errors were concentrated in a few slices in the most frontal and caudal positions. Since manual editing depends on both the average error and its distribution, such localized errors simplify manual correction when necessary.

### 3.2 Object detector accuracy

Two sources of error might affect SAMson’s accuracy: suboptimal prompt generation and model-specific errors that persist despite perfect bounding boxes. To evaluate these, we assessed the accuracy of our box generator (Fig. 4-a,b) and compared SAM’s oracle performance with SAMson’s Full-Auto performance (Fig. 4-c,d,e), determining the extent to which errors are attributable to our system versus inherent limitations of the segmentation model. For comparative analyses across datasets, we focused on the 3 m.o.a subgroup within the D1 dataset to minimize age-related variability, as it is closest in age to subjects in D2 and D3.

**Fig. 4:**
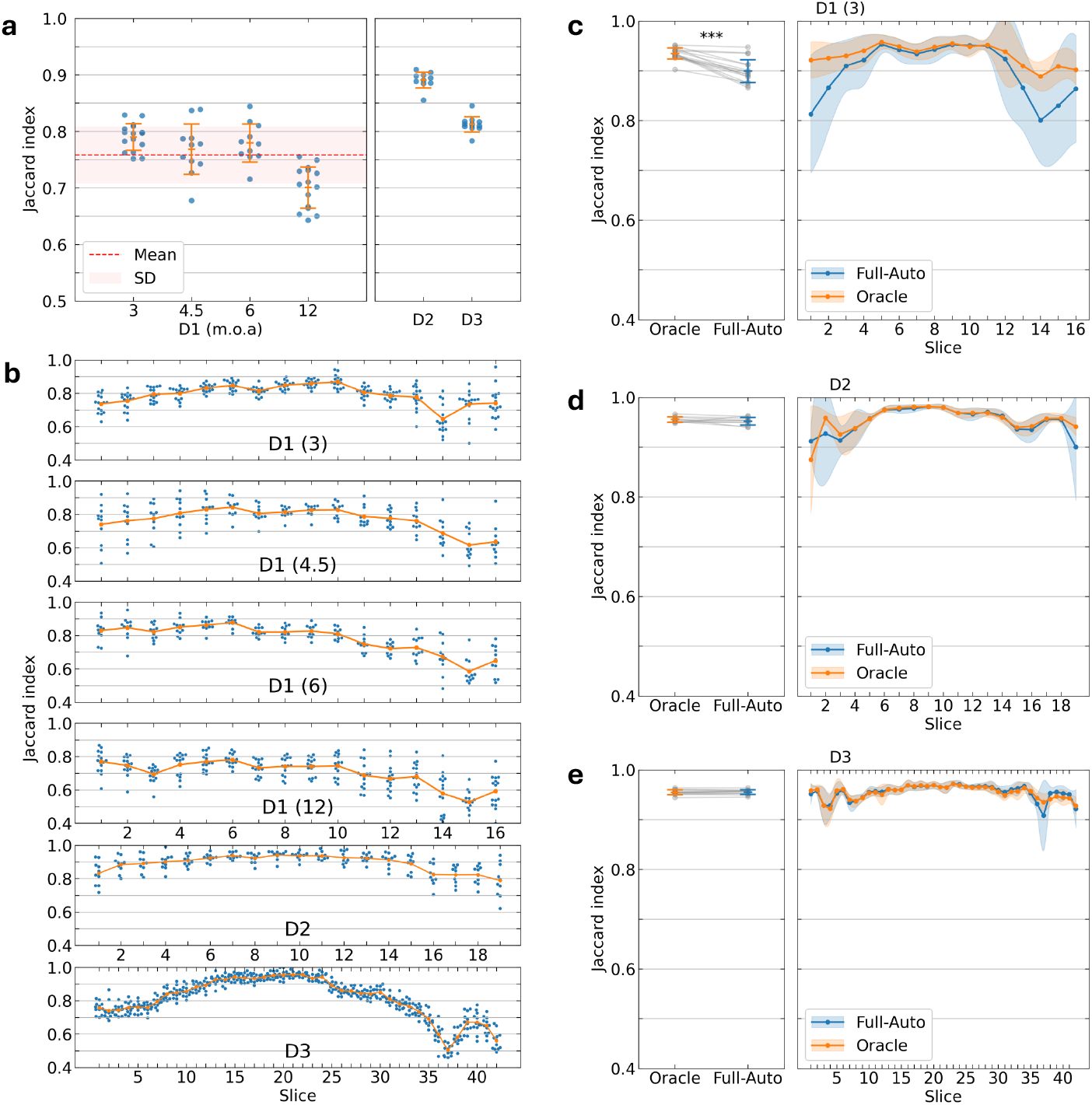
SAMson prompt generation accuracy and oracle performance. a,b) Accuracy analysis of all generated bounding boxes compared to GT boxes, defined as those that enclose GT masks. c-e) Comparison of SAMson’s oracle performance, achieved using GT boxes as input prompts, with its Full-Auto performance across datasets. Left: performance per subject, averaging across slices, where data points corresponding to the same subject are connected by grey lines. Paired t-test: c) *t*(30) = 5.31; *p* = 2.6 · 10^−5^, d) *t*(18) = 1.14; *p* = 0.27, e) *t*(22) = −0.14; *p* = 0.89. Right: performance per slice, averaging across subjects, with shaded areas indicating standard deviation.

For high-resolution images (D2 and D3), significantly more precise bounding boxes in D2 compared to D3 (unpaired t-test *t*(20) = 12.6; *p* = 1.22 · 10^−10^; Fig. 4a) did not result into significantly better mask outcomes (*t*(20) = 1.1; *p* = 0.27; Fig. 2a). In D3 images, the accuracy of bounding boxes across slices (Fig. 4b) showed moderate correlation (*r* = 0.43; *p* = 1.3 · 10^−23^) with the corresponding accuracy of mask generation using those boxes (Fig. 2b). For lower resolution images (D1 at 3 m.o.a), the variance of the object detector increased (Fig. 4a) and the correlation between suboptimal boxes and suboptimal masks also rose (*r* = 0.56; *p* = 9.0 · 10^−23^). This trend is reflected in the similarity of subject-age-wise and slice-wise trajectories for boxes and masks (Fig. 2-a,b and Fig. 4-a,b). Fisher z-Comparison showed that correlations between boxes and masks for D1(3) and D3 were significantly different (*z* = 2.36; *p* = 0.018), indicating that the impact of box prompt accuracy on mask generation quality becomes more pronounced as image resolution decreases.

The comparison between SAMson’s Full-Auto mode and the oracle performance showed no statistically significant differences in D2 and D3 datasets, either globally or across slices, (Fig. 4-c,d,e). However, a difference of small magnitude but statistically significant was observed in D1(3 m.o.a) dataset (paired t-test, Full-Auto vs. oracle performance, *t*(30) = 5.31; *p* = 2.6 · 10^−5^). This result indicates that, in most cases, the tool’s performance is not constrained by the accuracy of the object detector but by SAM’s capabilities in this specific MIS problem. Improvements to the object detector could enhance segmentation outcomes only for the most challenging slices (rostral and caudal) in low-resolution images, where the Semi-Auto mode already provides mask results nearly as precise as the oracle.

### 3.3 Comparison with other automated tools

To highlight the significance of the developed automated pipeline, we conducted a comparative analysis (Fig. 5) of its performance against established brain extraction methods, including BET [6] and deep learning-based tools such as RBM [10] and BEN [31]. For BET, we evaluated both the standard version optimized for mouse MRI data and BET4Animal, using the best-performing option. SAMson’s performance was compared across all three datasets, analyzing both mean accuracy per subject and per slice. SAMson consistently outperformed all methods under all tested conditions.

**Fig. 5:**
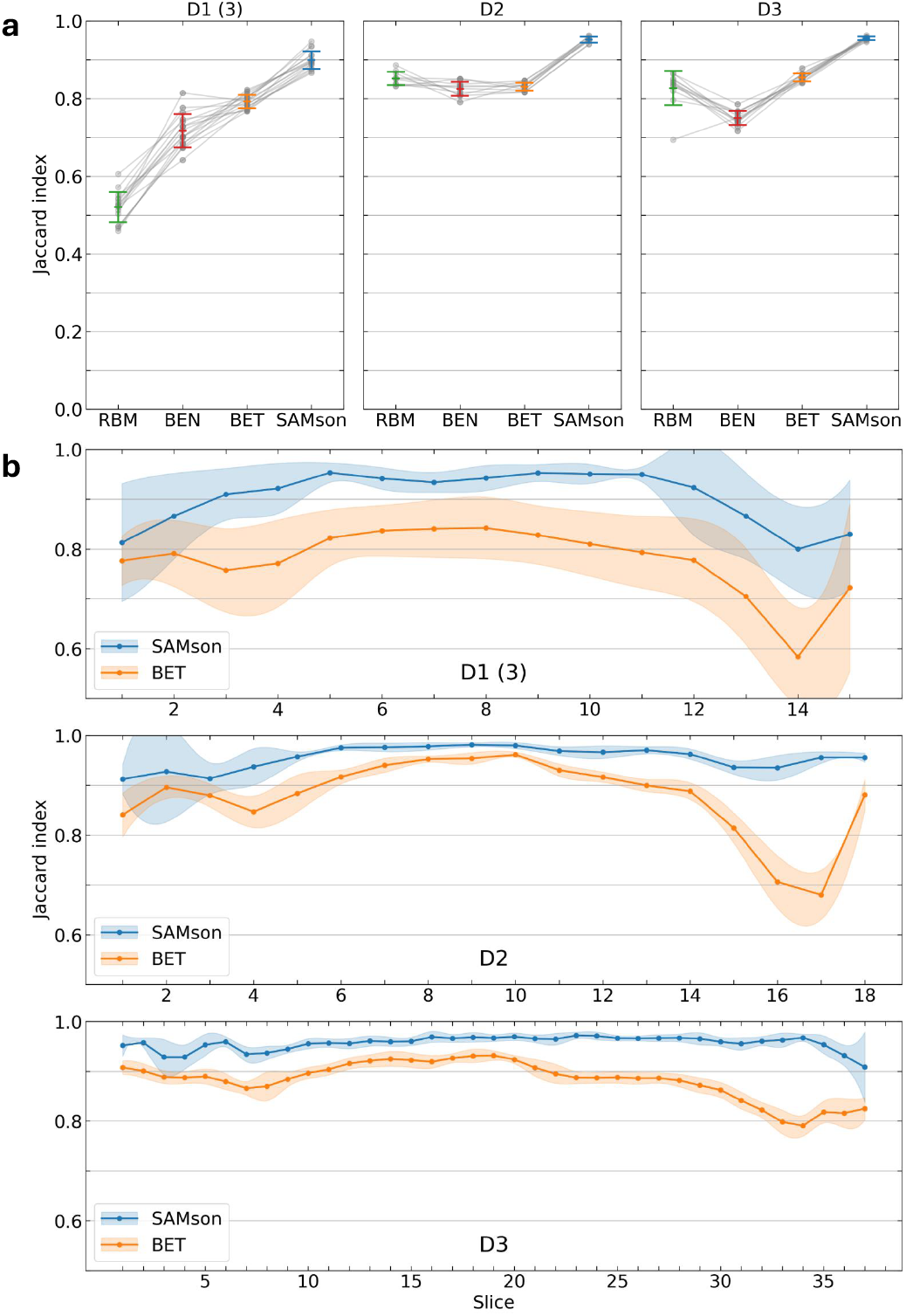
Benchmarking of RBM, BEN, BET and SAMson (Full-Auto). a) Average performance of each method across datasets, with results for the same subject from different tools connected by grey lines. b) Comparative analysis of SAMson and the second-best method, BET, based on their performance per slice, averaged across subjects for each dataset. Shaded areas indicate standard deviation.

## 4 Discussion

SAMson has demonstrated remarkable accuracy in segmenting rodent MRI brain images, outperforming currently available methods. While the quantitative analysis focused on T2-weighted images, the tool has also been successfully applied to other imaging modalities, such as diffusion-weighted MRI and fMRI. Its versatility is underscored by its two complementary approaches: Full-Auto mode provides efficiency and automation for most common applications and Semi-Auto is ideal for challenging scenarios.

The Semi-Auto mode requires the operator to select the most accurate mask between three options, which adds processing time but offers greater precision in challenging cases, such as low-resolution images. In these cases, SAM becomes more sensitive to prompt quality. Since low-resolution images typically contain fewer slices, the time required for manual selection might be less than the time needed to manually correct errors in fully-automated pipelines. As we depart from the ideal scenario, there is a compromise between image quality and the efficiency of both methods (Fig. 6).

**Fig. 6:**
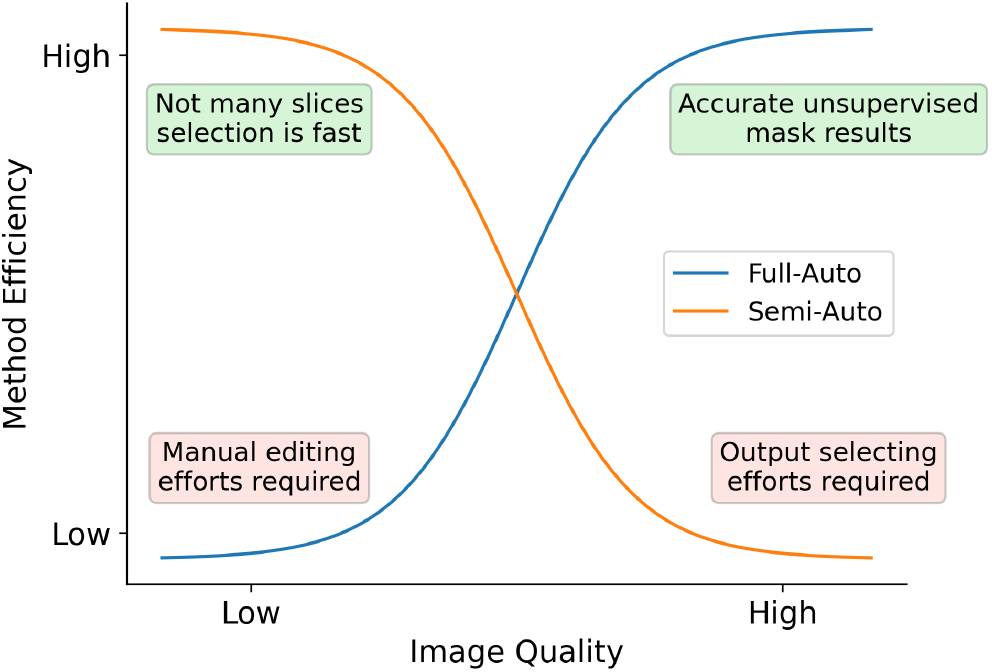
Compromise between image quality and SAMson method efficiency.

Automatically generated masks by SAMson rarely exhibit recurrent errors across slices, which can be attributed to two key factors. First, SAM’s tolerance to prompt inaccuracies allows it to adapt prediction beyond the specified bounding box. Second, the application of morphological transformations to SAM’s mask predictions consistently improves accuracy by removing false positive and marginally connected regions, enhancing alignment with the nuances of our specific segmentation problem.

It is noted that SAMson’s errors tend to be localized within specific (most rostral and caudal) slices, facilitating mask refinement when necessary and enhancing operational efficiency. This can be attributed to SAM’s 2D slice-based segmentation approach, which generates independent predictions for each slice. In contrast, volumetric brain extraction tools, such as BET, propagate errors across slices, compounding inaccuracies. The rostral and caudal localization of errors probably reflects partial volume effects—due to mixed brain and extracerebral tissue in the same voxel—limiting tissue boundary definition. Improved performance on higher-resolution datasets supports this interpretation. Also, since the Jaccard index measures relative overlap, it is particularly sensitive to differences in mask size. Smaller masks, such as the rostral and caudal ones, experience a disproportionately larger relative decrease in overlap for any deviation from GT.

The reduced performance of D1(12 m.o.a) compared to younger subjects likely stems from limitations in the study’s template, which was derived from younger animals and may not represent older anatomical features. This resulted in lower bounding box accuracy in aged animals, which as we have shown translated into segmentation mistakes. Future extensions of SAMson will include age-template selection for a particular study or group.

In the omics era, the preprocessing of large datasets requires automation for scalability and efficiency. SAMson streamlines brain extraction, significantly reducing manual effort while maintaining high accuracy. Its adaptability across various rodent experiments, scanner resolutions, and imaging conditions highlights its utility. Future iterations will focus on enhancing segmentation performance and expanding functionality to other preclinical species. By offering open-source code and data, we aim to enable the scientific community to adopt and improve SAMson, advancing neuroscience research.

## CRediT author statement

D.P.S.: Conceptualization, Software, Data curation, Resources, Formal analysis, Validation, Investigation, Visualization, Methodology, Writing – original draft, Writing – review & editing. M.K.S.: Conceptualization, Methodology, Writing – original draft. P.M-T.: Data curation, Resources. E.M-M.: Resources. P.R-C.: Resources. P.L-L.: Resources. S.D.S.: Conceptualization, Resources, Supervision, Project administration, Writing - original draft. S.C.: Conceptualization, Resources, Supervision, Funding acquisition, Project administration, Writing – original draft, Writing – review & editing. A.P.: Conceptualization, Supervision, Project administration, Writing - original draft, Writing - review & editing.

## Acknowledgements

S.C. acknowledges support from PCI2024-153491 funded by MICIU /AEI /10.130 39/501100011033 and UE, PID2021-128158NB-C21 funded by MICIU/10.13039/ 50110 0011033 and FEDER, UE, CEX2021-001165-S funded by MICIU/AEI /10.13039/50110 0011033 and Excellence Grant CIPROM/2022/15 and INVESTIGO INVEST/2022/396 funded by the Generalitat Valenciana and Next Generation EU. S.D.S. was supported by the the Spanish Ministerio de Ciencia e Innovación, Agencia Estatal de Investigación (PID2021-128909NA-I00 and CNS2023-14488), by the Programs for Centres of Excellence in R&D Severo Ochoa (CEX2021-001165-S), and by the Generalitat Valenciana through a Subvencion para la contratación de investigadoras e investigadores doctores de excelencia 2021 (CIDEGENT/2021/015) and by a Fundación Pasqual Maragall Research Programme (2024 call). P.L-L. was funded by grant PID2021-122528OB-I00 funded by MICIU/AEI/10.13039/501100011033/FEDER, UE. Some of the mice used in this study were generated in the project funded by MCIU/AEI/FEDER, UE grants PID2022-141700OB-I00, MCIN/AEI/10.13039/501100011033 and CB/07/09/0034 Center for Networked Biomedical Research on Mental Health (CIBERSAM, Bortolozzi A). D.P.S. gratefully acknowledges Gaspar Panadero Tendero for inspiring the initial idea of this work.

## A Algorithms

### Algorithm 1

Box Prompt Generation

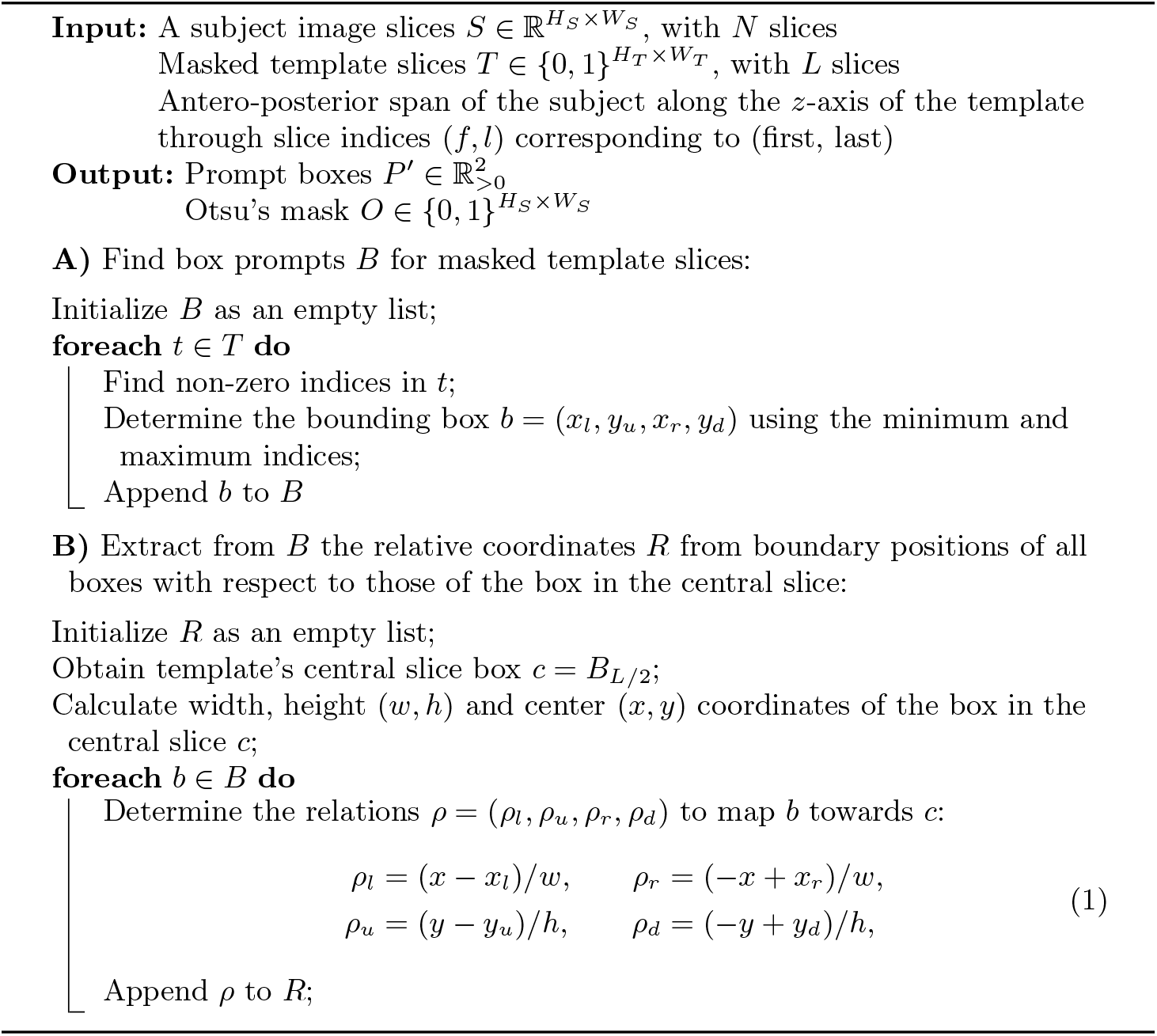
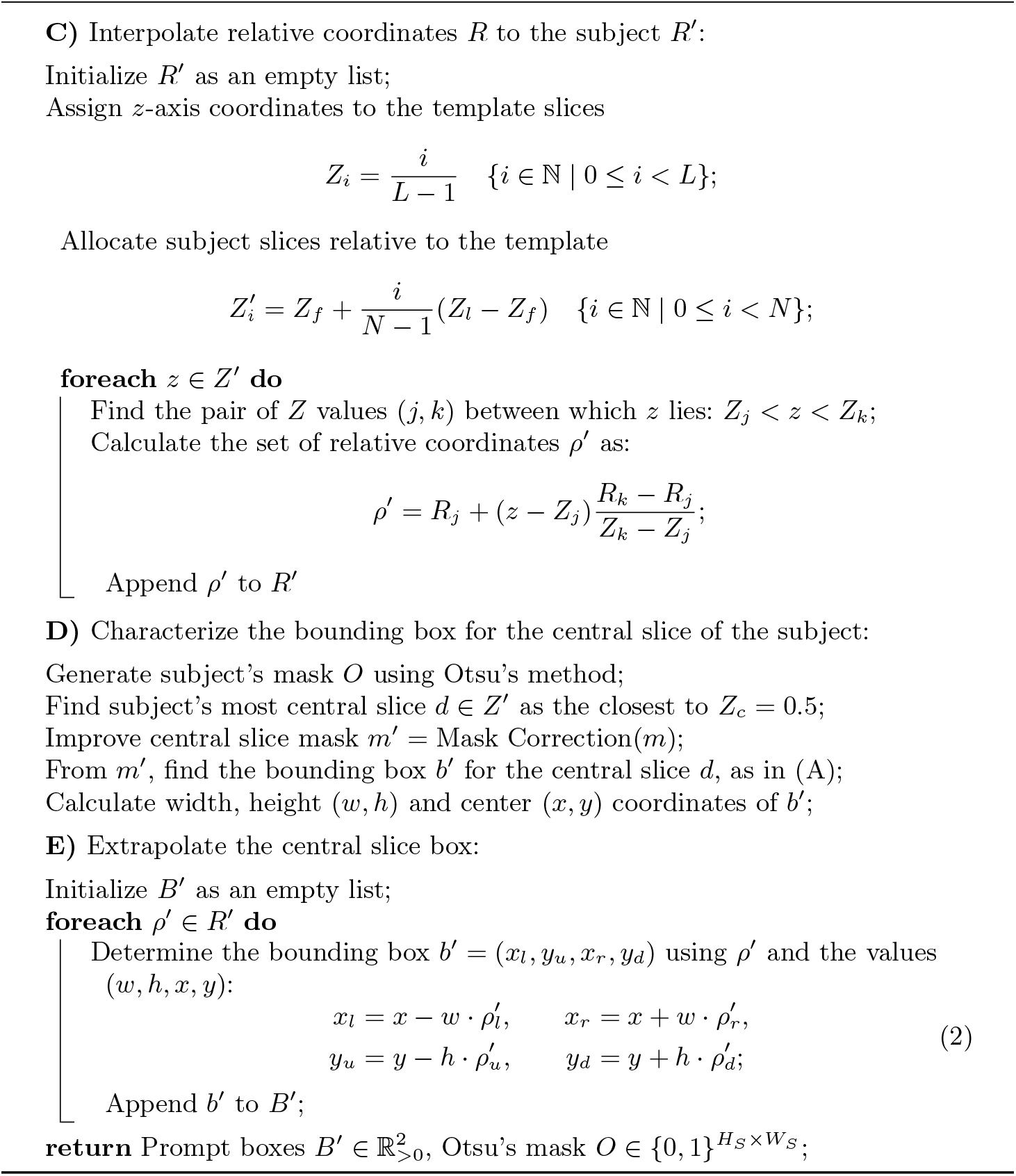

### Algorithm 2

SAMson Semi-Auto Mask Generation

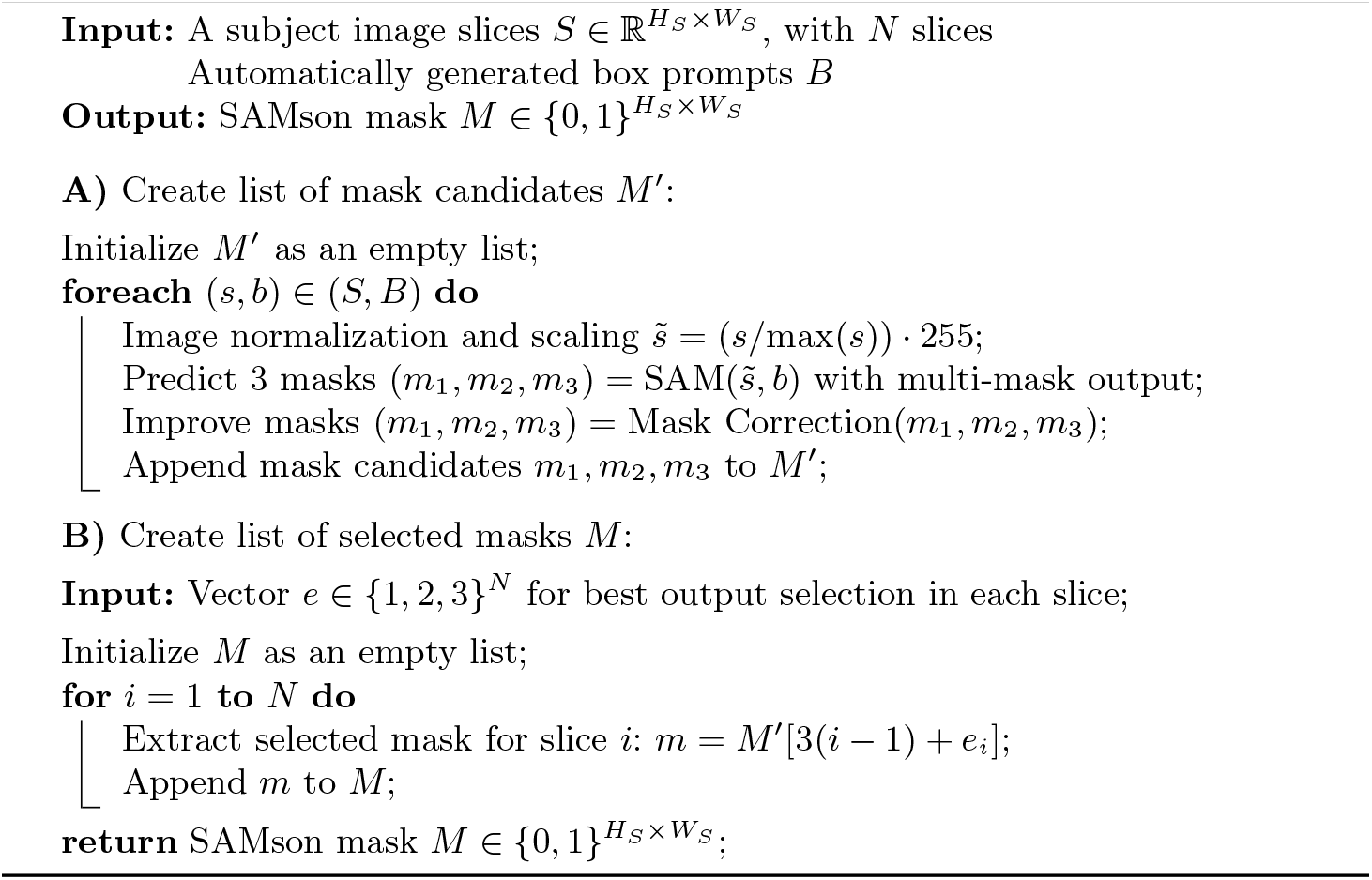

### Algorithm 3

SAMson Full-Auto Mask Generation

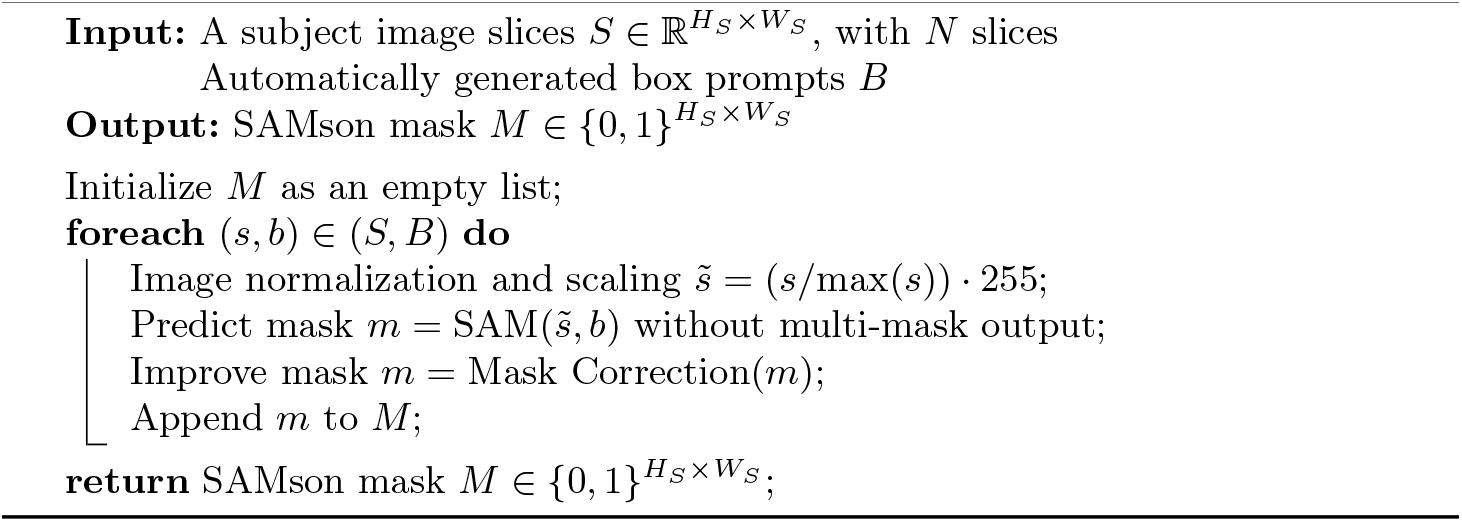

### Algorithm 4

Mask Correction

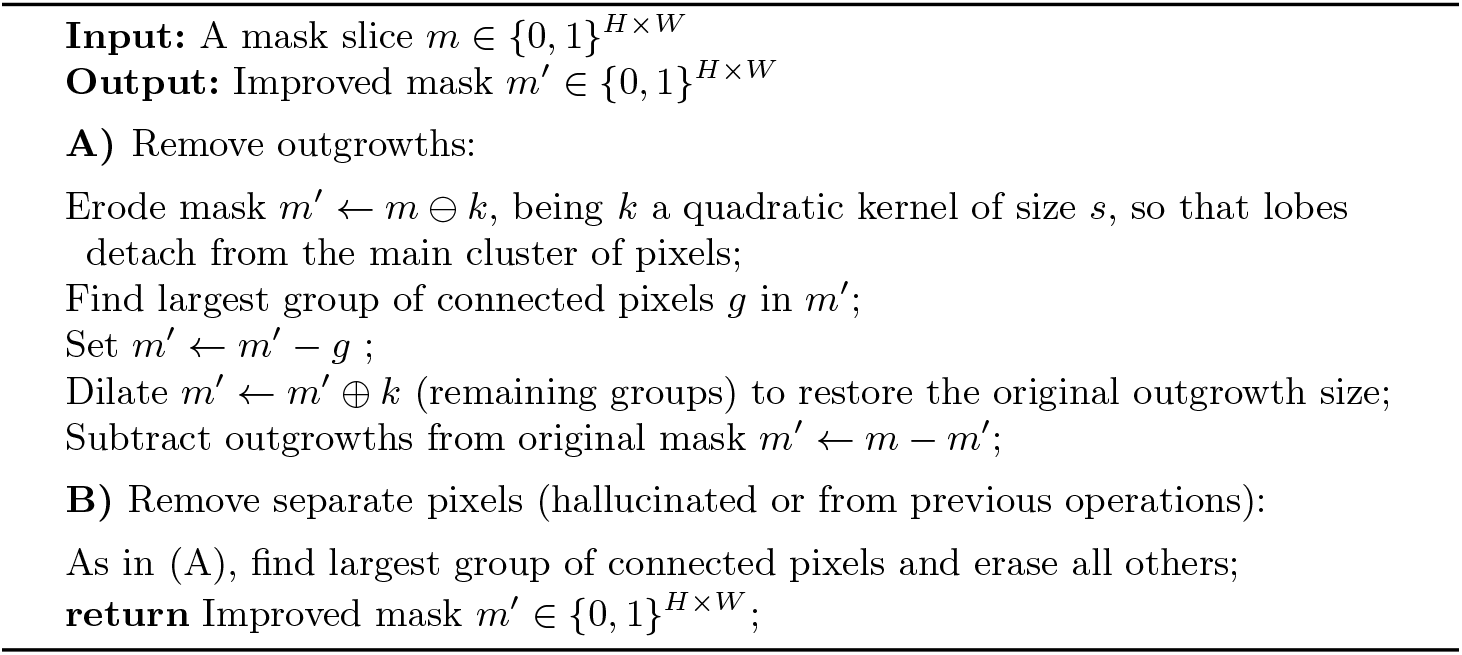

## Footnotes

1 https://github.com/facebookresearch/segment-anything

